# Self-Organization of Spindle-Like Microtubule Structures

**DOI:** 10.1101/624874

**Authors:** Bianca Edozie, Sumon Sahu, Miranda Pitta, Anthony Englert, Carline Fermino do Rosario, Jennifer L. Ross

**Author notes:** Corresponding author to whom correspondence should be sent.

## Abstract

Microtubule self-organization is an important physical process underlying a number of essential cellular functions including cell division. In cell division, the dominant organization is the mitotic spindle, a football-shaped microtubule organization. We are interested in the underlying fundamental principles behind the self-organization of the spindle shape. Prior biological works have hypothesized that motor proteins are required for the proper formation of the spindle. Many of these motor proteins are also microtubule-crosslinkers, so it is unclear if the important aspect is the motor activity or the crosslinking. In this study, we seek to address this question by examining the self-organization of microtubules using crosslinkers alone. We use a minimal system composed of tubulin, an antiparallel microtubule-crosslinking protein, and a crowding agent to explore the phase space of organizations as a function of tubulin and crosslinker concentration. We find that the concentration of the antiparallel crosslinker, MAP65, has a significant effect on the organization and resulted in spindle-like organizations at relatively low concentration without the need for motor activity. Surprisingly, the length of the microtubules only moderately affects the equilibrium organization. We characterize both the shape and dynamics of these spindle-like organizations. We find that they are birefringent homogeneous tactoids. The microtubules have slow mobility, but the crosslinkers have fast mobility within the tactoids. These structures represent a first step in the recapitulation of self-organized spindles of microtubules that can be used as initial structures for further biophysical and active matter studies relevant to the biological process of cell division.

## Introduction

Biological materials and structures have the amazing capacity to self-organize into useful architectures that can perform productive work at a multitude of length and time scales. From protein assemblies to whole animals, biological systems are machines made up of smaller machines. These machines are capable of building, repairing, and annihilating themselves using active, energy-using processes.

One particularly intriguing system that has these properties is the mitotic spindle of the dividing cell. The spindle is a self-organized network of filaments that undergoes the task of aligning and separating genetic material (DNA in chromosomes) of cells into two new daughter cells, an essential process in biology. Indeed, self-replication is often the defining feature of “life.” Yet, we still do not fully understand how the mitotic spindle is formed, organized, repaired, or broken down.

Microtubule cytoskeletal filaments are Integral to the process of cell division. Microtubules are rigid, hollow rod filaments composed of tubulin dimers. The dimers are enzymes that use the energy of GTP hydrolysis to self-immolate. Their polymerization is driven by entropy and the desire of water molecules to be away from hydrophobic surfaces. GTP hydrolysis by the tubulin enzyme changes the dimer shape and interactions to cause the filament to disassemble in an unzipping wave from their ends (for recent reviews, see ^1–3^).

The self-organization of microtubules into higher order structures has been the subject of interest for over 20 years when it was demonstrated that microtubules can form patterns of asters ^4,5^. Recent advances using microtubules as active liquid crystals and self-organized shuttles ^6–11^ have used the power of associated motor proteins to rearrange microtubules. However, these past experiments did not take advantage of microtubules’ inherent polymerization machinery. A more recent result created an exquisitely complex system of dynamic microtubules and crosslinking motor proteins that could dial the organization of the microtubules from astral for slowly polymerizing microtubules to nematic for quickly polymerizing microtubules ^12^, demonstrating that the polymerization rate is an important knob for microtubule self-organization.

Here, we seek to create a minimal system of self-organizing microtubules using the polymerization of the filaments as the driving propulsion and crowding agents to enhance density. We increase the complexity by adding adhesion between the typically repulsive filaments, using a crosslinking protein, MAP65. MAP65 is an antiparallel crosslinker found in plants that is analogous to PRC1 from animals and Ase1 from yeast. We have previously shown that MAP65 can organize gliding microtubules into active structures that are reminiscent of cellular architectures in dividing cells ^13,14^. MAP65, Ase1, and PRC1 have each been shown to be weakly-associating crosslinkers that can drive microtubule bundling in vitro ^15–18^.

Using this simple, yet powerful, experimental system, we show that we can create spindle-like assemblies of microtubules that nucleate and grow until they reach equilibrium using crosslinking elements alone. Without the crosslinker, the filaments locally align into ordered domains. With increasing adhesion, the assemblies condense into spindle-like structures. We characterize these structures and demonstrate that their shape is reminiscent of the mitotic spindle, yet they are not liquid-like. Our results demonstrate that the spindle-like architecture found in dividing cells only requires crosslinking, but not motor activity.

## Experimental

### Chemical Reagents

All reagents were purchased from Sigma Aldrich unless otherwise specified.

### Proteins

Proteins used in this study include tubulin and MAP65. Tubulin purified from pig brains was purchased from Cytoskeleton (Denver, CO) as unlabeled or fluorescently labeled, lyophilized protein. Lyophilized tubulin was hydrated with PEM-80 (80 mM PIPES, pH 6.8, 1 mM MgSO_4_, 1 mM EGTA) and incubated for 10 minutes on ice. Hydrated tubulin was centrifuged at 366,000 × g for 10 minutes at 4°C to remove aggregated tubulin. Tubulin was kept on ice until insertion into chamber for experiments.

MAP65-1 plasmid with or without GFP was a gift from Ram Dixit (Washington University, St. Louis). MAP65 was expressed and purified from bacteria, as previously described ^13,14,19^. Briefly, bacteria were transformed with the plasmid and grown to an OD of 1. Soluble protein was harvested from bacteria through centrifugation and affinity purified using the 6xHis tag binding to nickel ion. Protein was stored at -80°C.

### Experimental Chamber

Assays were carried out in 10 μL flow chambers made of a glass slide attached to a coverslip with permanent double-stick tape (3M). Before use, coverslips were cleaned with ethanol and water. Clean coverslips were treated with dimethyldichlorosilane solution, 2% (wt/vol) (GE Healthcare) as previously described ^20,21^ to create a hydrophobic surface that was blocked with Pluronic F127 block-copolymer in experiments.

### Experimental Procedures

Microtubule lengths were measured using the following procedure. Unlabeled and fluorescently labeled (rhodamine), lyophilized tubulin was hydrated with PEM-80 to the concentration required. The rhodamine tubulin was mixed with the unlabeled tubulin at a 1:10 ratio and allowed to incubate on ice for 10 minutes. The tubulin was centrifuged for 10 minutes at 360,000 × g in 4°C to remove aggregated tubulin. The supernatant was transferred to a new tube, GMPCPP (Jena Biosciences) was added to a final concentration of 0.6 - 3 mM, and the mixture incubated at 37°C for 30 minutes to polymerize the filaments. After polymerization, the microtubules were diluted 100x into new PEM-80, added to a slide and coverslip, and imaged in epi-fluorescence. There was no significant difference between filaments at 0.6 mM and 3 mM GMPCPP.

Microtubule self-organization experiments were performed using the following procedure. Unlabeled and fluorescently labeled (Rhodamine or Dylight 488), lyophilized tubulin was hydrated with PEM-80 to the following stock concentrations: 54.5 μM or 182 μM and diluted from there or drop frozen for future use. Tubulin was diluted to the working concentration (13.6μM, 54.5 μM, 70 μM) incubated in buffer for 10 minutes at 4°C and then mixed at a 1:10 ratio, labeled to unlabeled. The tubulin was centrifuged at 360,000 × g for 10 minutes in 4°C to remove aggregates, and kept on ice until ready to be mixed and polymerized in the chamber. While the tubulin was being prepared, the experimental chamber was assembled from the silanized coverslip, slide, and double stick tape. A surface blocking agent, 5% Pluronic F127 block copolymer, was added to the chamber and incubated for 5-10 minutes. Prior to insertion into the chamber, the tubulin was mixed with the following: 3 mM GMPCPP to polymerize and stabilize microtubules, 0.5% Pluronic F127 to maintain surface coating, 50 mM DTT as a reducing agent to inhibit disulfide bonds and scavenge oxygen, 15 mg/mL glucose, 0.5 mg/mL glucose oxidase, 0.15 mg/mL catalase as an oxygen scavenging system, 0.3% methylcellulose (4,000 cP) as a crowding agent, MAP65 in PEM-80 to correct concentration as the crosslinker, and tubulin at the correct concentration to polymerize into microtubule organizations inside the chamber. The sample chamber sealed with epoxy (Z-poxy) to prevent dehydration and incubated at 37°C in a closed, temperature controlled incubator to polymerize the filaments for 30 min – 4 hours. The concentration of MAP65 added depended on the concentration of tubulin, the desired percent bound, and the equilibrium dissociation constant for MAP65 binding to microtubules, K_D_ = 1.2 μM ^19^. A full calculation and list of the exact concentrations used in this study can be found in the supplemental methods (ESI Table 1).

### Imaging

Microtubule lengths were imaged by epi-fluorescence microscopy using a Nikon Ti-E microscope with Perfect Focus, 60x oil (1.49 NA) or 60x water (1.20 NA) immersion objectives, and illuminated with a XeHg lamp or LED lamp system. Nikon Elements was used to record images from the camera, either an S-CMOS (2048×2048 pixels) Zyla (Andor) or an electron multiplier CCD (512×512 pixels) Ixon (Andor). The pixel size for the CMOS was 108 nm/pixel and for the CCD was 108 nm/pixel after including a 2.5x lens in front of the camera. Rhodamine fluorescence was imaged with a standard dichroic filter set for TRITC (ex: 540, em: 605, Chroma).

Microtubule self-organizations in the presence of crowders with or without MAP65 were imaged using total internal reflection fluorescence (TIRF) microscopy. A Nikon Ti-E microscope with Perfect focus and 60x oil (1.49 NA) immersion lens with extra 2.5x magnification before the Andor Ixon EM-CCD camera to give a 108 nm/pixel. TIRF illumination was performed with a custom-built laser combination and manipulation system that coupled to the epi-illumination path in the back of the microscope ^22^. For dynamic imaging experiments, we warmed the tubulin sample to 37°C using an air heater to blow onto the sample. The final state of experiments performed while imaging were identical to those prepared in the incubator, as described above. We used Nikon Elements software to record image data and time series data. For dynamics of assembly measurements, images were recorded every 5 sec for 2-4 hours. For samples incubated off the microscope, at least 10 individual images were taken throughout the chamber.

Fluorescence recovery after photobleaching (FRAP) was performed using a Nikon Ti-E microscope with Perfect Focus and 60x oil immersion (1.49 NA) objective imaged with an EM-CCD Ixon camera (Andor) in the UMass Institute for Applied Life Science light microscopy user facility. Spinning disc imaging was performing using a Yokogawa Spinning Disk Confocal using 488 nm laser for imaging GFP, 638 nm laser for imaging DyLite-650 tubulin or 532 nm laser to image rhodamine-tubulin. FRAP was performed with Orthogonal Stimulation using 405 nm to bleach both green (GFP-MAP65) and red (tubulin) at the same time. After 60 seconds of imaging one frame per 2 seconds, the region was photobleached and imaged for 30 minutes.

### Data Analysis and Statistics

Microtubule lengths, tactoid lengths, and tactoid widths were measured from still images of microtubules using FIJI/ImageJ by hand tracking individual filaments. Data was converted to length in micrometers using the pixel size conversion.

Orientation domain areas were identified and measured using the OrientationJ plugin in FIJI/ImageJ. The Gaussian window was set to 5 pixels using the cubic spline gradient setting. The color based on orientation and brightness was set either using the intensity of the original image to show the filaments, or constant, to show the domains only. In the image of the domains (constant brightness), the color threshold was used to select regions of the same color range. The total color range was from 0 to 255, and the selected ranges were 0 to 63, 64 to 127, 128 to 191, and 192 to 255. When a color range was selected, the thresholded region was colored white and selected using the magic wand tool. Once selected, we measured the number of pixels within the selected area. Regions may be divided into two orientation domains if the filaments within the region have angles/colors that are close to the cut-off between color ranges listed above. The selection and measurement was performed on each region within the image. We used 6 different images from 3-4 different chambers to make the measurement.

Tactoid shape was quantified by rotating all tactoids to have their long axes horizontal, thresholding the image to be black-and-white, removing other shapes smaller than our interest in the background, and then measuring the location of edge of the profile using MATLAB edge detection algorithm. The distance between the two edges of the profile were denoted as the width, which was recorded for each pixel location along the length. The length (length of the array) and maximum width (maximum value of that array) of the tactoid were measured and used to rescale the data. Because each tactoid was a slightly different length, and we sought to average the data, we took 20 measurements along the length by averaging the widths over these 20 bins. After averaging over 10 tactoids, the mean and standard deviation for each bin was plotted as the rescaled width as a function of the rescaled length. Data was fit with a parabola as given in the results and best fit parameters are given in the ESI.

FRAP was measured from time series data recorded using spinning disc microscopy. The photobleached region was selected and the integrated intensity was measured within the region for all frames. To account for background photobleaching, we also measured the same sized region in a location that was away from the photobleached region. The background displayed a linear photobleaching that was corrected with a linear offset prior to analysis for photobleaching. Fluorescence intensity data was rescaled for each location between 0 and 1 by measuring the average intensity over the first 1 minute of recording when there was no intentional photobleaching, and setting that to the initial intensity. The intensity of the first bleached frame was measured, and the difference between the initial and minimum intensity was computed. The integrated intensity was rescaled by first subtracting the minimum intensity from each frame and then dividing by the difference between the minimum and the initial intensity. Recovery data of this type was fit to an rising exponential decay function. Kymographs of the intensity during the FRAP experiment were created using the MultiKymograph plugin in FIJI/ImageJ. The intensity along the tactoid was measured for specific time points using the kymograph.

### Results and discussion

The process of polymerization is a driving force in active self-organization processes found in live cells. We are interested in the ability of microtubules to self-organize during polymerization from dimers into filaments (Fig. 1A). We have also added an additional component to cause crosslinking during the process, MAP65, a plant-derived, microtubule-crosslinker that preferentially bundles microtubules in antiparallel organizations (Fig. 1A). When combined in the experimental chamber, free tubulin dimers nucleate and grow filaments that can quickly crosslink via the MAP65 to form small assemblies of microtubules (Fig. 1A). These assemblies can grow in length, through the addition of free dimers to the ends of microtubules, and in width, through the association of new microtubules to the existing bundle (Fig. 1A). The process can be observed over time to quantify the dynamics of the assembly and characterize the final steady state with or without crosslinking proteins (Fig. 1B).

**Figure 1.**
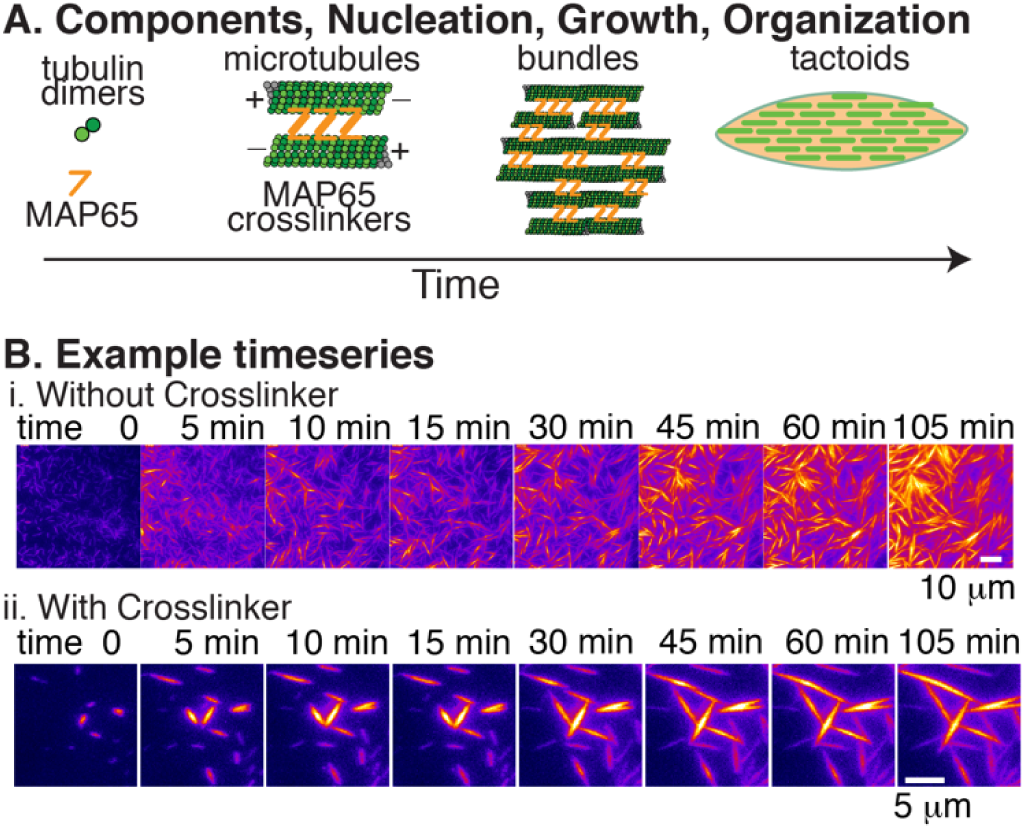
Experimental design. (A) Components that nucleate and grow to self-organize during experiment include tubulin dimers (green) that self-assemble into microtubules and the antiparallel microtubule-crosslinking protein, MAP65 (orange). Microtubules crosslink into bundles that form tactoids. (B) Timeseries examples of self-assembly and organization (i) without crosslinkers and (ii) with crosslinker. (i) Sample shown with 13.6 μM tubulin in the absence of MAP65. Time between frames is 5 min; scale bar is 10 μm. (ii) Sample shown with 13.6 μM tubulin in the presence of MAP65 where 3 tubulin dimers have a MAP65 bound for each 200 dimers (1.5%). Time between frames is 5 min; scale bar is 5 μm.

### Microtubule length control

When assembling structures from filaments, the length distribution of the filaments may have a large impact on the resulting structure. Specifically, it is known that the aspect ratio of liquid crystal molecules can affect the transition to various phases ^23^. Biology seemingly uses this length control to facilitate both microtubule and actin network architectures. Prior work with actin is facilitated by the fact that actin has a number of known associated proteins that can control the filament length ^24–26^. Microtubule associated proteins that can help control length do so indirectly, which is distinct from actin binding proteins that can cap to directly control the length of filamentous actin.

Since microtubules do not have easy length-control proteins, we controlled polymer length through controlling the nucleation and growth kinetics, which are both controlled by the concentration of the filaments. Unfortunately, microtubules undergo a process called dynamic instability where individual filaments grow and stochastically switch to a shrinking phase. This switch from a growing phase to a shrinking phase is caused by a conformational change upon hydrolysis of the GTP molecules to a GDP molecules (for several recent reviews, please see ^1–3^. The shift to depolymerization destabilizes the nuclei that initially form, resulting in fewer, longer filaments.

To avoid the effects of dynamic instability, microtubules can be stabilized through small molecules, such as paclitaxel (Taxol) or a slowly-hydrolyzable analog of GTP such as GMPCPP. These small molecules reduce the critical concentration for nucleation and stabilize the filaments that polymerize. Stabilization results in many, shorter filaments, and allows length control as a function of initial dimer concentration. Specifically, higher concentrations increase the filament nucleation rate to result in more, shorter filaments. This method was previously employed to control the length of microtubules for active liquid crystal measurements ^27^.

Since each lab may have different results for microtubule length control, we quantified the length distribution as a function of initial tubulin concentration in our hands (Fig. 2). We find an empirical relationship between the filament length and initial tubulin concentration by polymerizing tubulin in the presence of GMPCPP at five different tubulin concentrations: 13.6 μM, 15.5 μM, 45.5 μM, 46.4 μM, and 54.5 μM in separate containers at 37°C. As a control to typical polymerization and growth conditions, we also polymerized microtubules using our typical assembly methods: polymerizing 45.5 μM in the presence of 1 mM GTP at 37°C and stabilizing with Taxol. After polymerization, the filaments were diluted to 0.5-1.0 µM and inspected via fluorescence microscopy. Dilution should not alter the length since the filaments are all stabilized.

**Figure 2.**
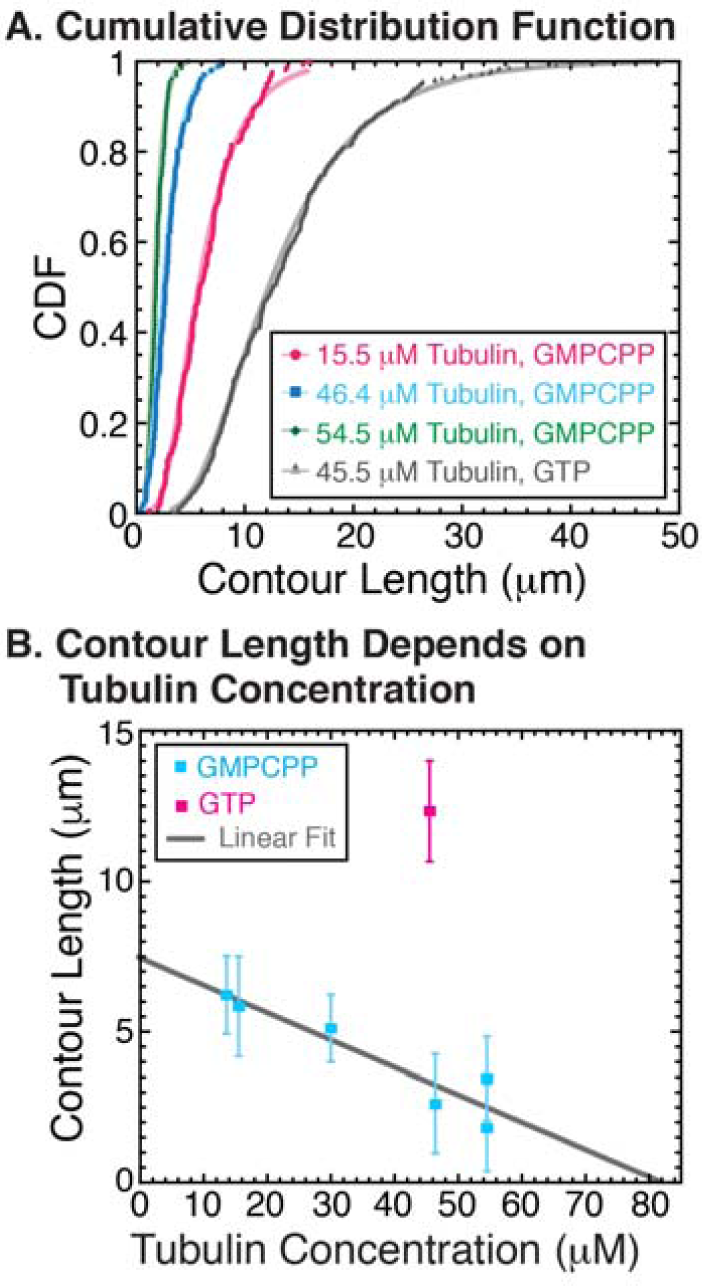
Microtubule length control. (A) Three concentrations of tubulin dimers were polymerized in the presence of GMPCPP: 15.5 μM (n = 162, pink), 46.4 μM (n = 101, blue), 54.5 μM (n = 127, green), and compared to 45.5 μM tubulin polymerized in the presence of GTP (n = 403, gray). Cumulative distribution functions (CDF) of lengths were lognormally distributed for all experiments, as expected for nucleation and growth polymer kinetics. CDFs were fit to Eq 2; fit parameters are given (ESI Table S2). (B) Using the medians determined from the fits to the CDFs (squares), and plotting as a function of tubulin concentration revealed a linear dependence on microtubule length with GMPCPP (blue squares, gray fit line, ESI Table S3). Microtubules made with GTP (pink squares) are outside the linear dependence.

Cumulative probability distributions of the measured filament lengths displayed lognormal distributions, typical for polymerization kinetics (Fig. 2A). From the plotted data, it was apparent that the median length decreased as a function of increasing tubulin concentration in the presence of GMPCPP. In addition, the width of the distribution decreased with increasing tubulin, making the length population more uniform. The median lengths and distribution widths were largest for the control microtubules polymerized with GTP and stabilized with Taxol (Fig. 2A).

We plotted and fit the cumulative distribution function (CDF) data because the probability distribution functions are prone to uncertainty due to binning the data. CDFs were fit to the expression for a lognormal distribution, with the form:

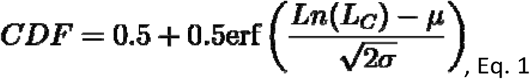

where the *µ* and *σ* are the mean and standard deviation of the normal distribution of *ln(x)* (Fig. 2A). The fit provides information about the most common length as a function for each concentration of tubulin. To obtain the characteristic length, we compute the exponential of *µ* from each fit to find the median contour length. We plotted the median *L*_*C*_ as a function of the initial tubulin concentration, and the data fit well to a line with negative slope for data polymerized with GMPCPP (Fig. 2B, fit parameters given in ESI Table S3). The typical Taxol stabilized microtubules do not scale with the GMPCPP data because these filaments were able to undergo dynamic instability prior to stabilization with Taxol, resulting in fewer, longer filaments. Using the fit results, we estimate the median contour length of microtubules as a function of the initial tubulin concentration for all the following data.

### Microtubule organization without crosslinking

Using the empirically derived length dependence, we chose to examine the microtubule self-organization during polymerization using three different free tubulin concentrations of 13.6 μM, 54.5 μM, or 70 μM which should result in median lengths of 6.2, 2.0, or 1.1 μm, respectively. All experiments were performed in the presence of GMPCPP. We also included 0.3% methylcellulose to act as a crowding agent to force the filaments to the cover glass surface as they organized. This served to reduce the dimensionality of the experiment and enabled imaging using total internal reflection fluorescence (TIRF) microscopy. The cover glass was passivated with a block copolymer, Pluronic F127, to keep tubulin dimers from sticking to the surface during the experiment.

When microtubules are grown without crosslinker, with only methylcellulose present, they form patterns that are locally aligned, but isotropic on larger scales (Fig. 3A). The common motif is fan-like splayed organizations of bundles of filaments (Fig. 3A). Since methylcellulose is a crowding agent, it will not only enhance the filament interaction with the surface, but will also help to bundle the filaments. This has been previously observed with poly-ethylene glycol and dextrans ^7,28,29^. Timeseries data capturing the nucleation and growth show many microtubule bundles nucleating and growing throughout the surface (Fig. 1B). Filament bundles have some mobility until they grow into each other and become locally jammed into the fan-like patterns. The pattern of local alignment arrests fairly early in the process to cause splaying organizations of filaments throughout the chamber (Fig. 3A).

**Figure 3.**
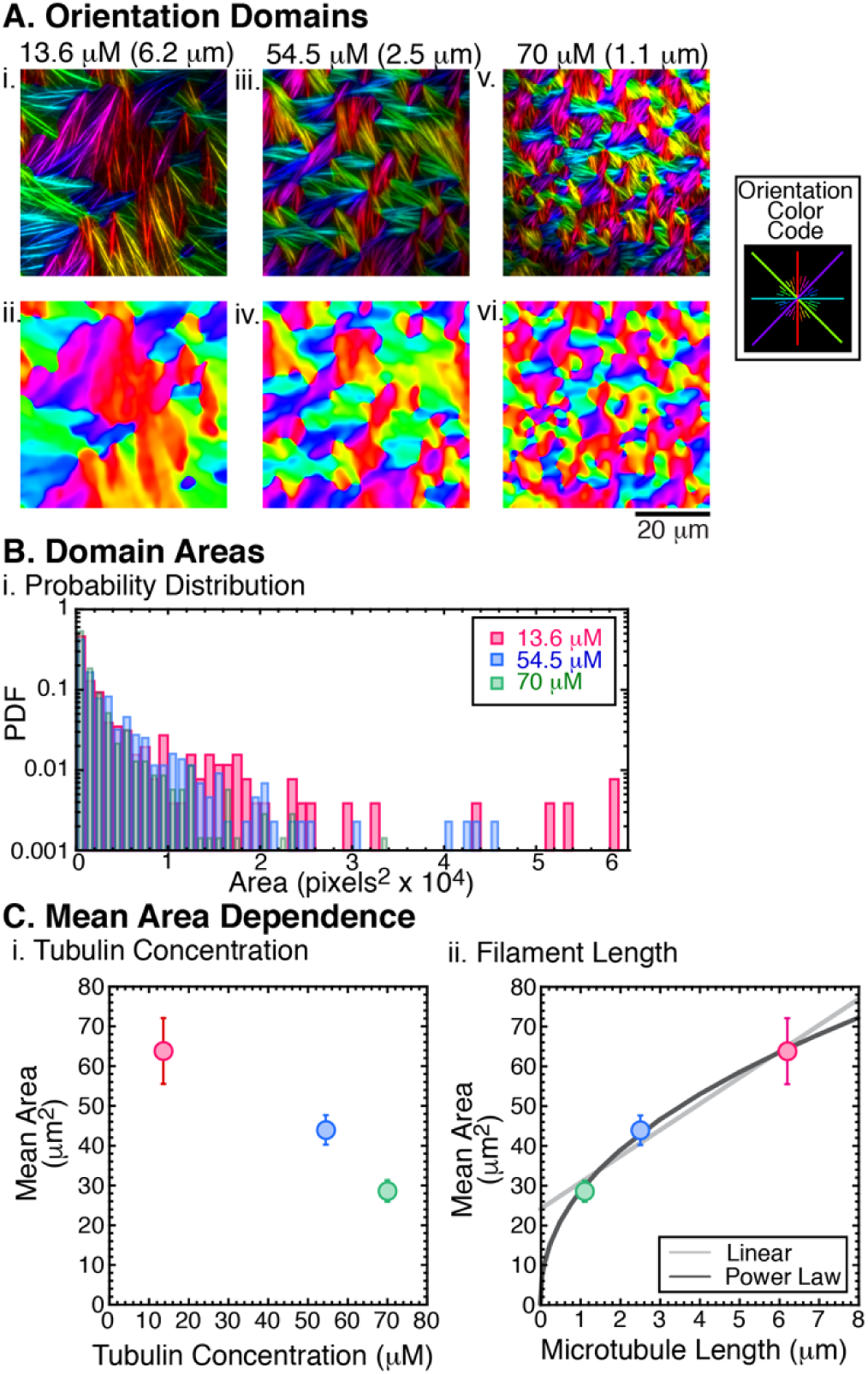
Orientation Analysis. (A) Polymerization of tubulin at (i,ii) 13.6 μM, (iii, iv) 54.5 μM, and (v, vi) 70 μM in the absence of crosslinkers created fan-like organizations of microtubules on the surface displayed with the filaments (i, iii, v) or without the filaments (ii, iv, vi). Color coding based on orientation displayed in legend. Scale bar is 20 μm for all images. (B) Areas of the color domains from (A) plotted as probability distribution histograms for 13.6 μM (n = 260, pink bars), 54.5 μM (n = 437, blue bars), and 70 μM (n = 703, green bars). (C) The mean area of the orientation domains plotted as a function of (i) tubulin concentration and (ii) filament length. Area vs. Length was fit to linear (light gray) and power law (dark gray) equations. Fit parameters are given (ESI Table S4). Error bars represent the confidence interval in the mean.

Using the OrientationJ plugin in FIJI/ImageJ, we automatically determine the local orientation of the filaments and color coded the director field (Fig. 3A). The area of each region pointing in a specified direction is determined by thresholding on a range of colors, selecting the region, and measuring the area within the domain in pixels. One region could be split into two, if the angle was at the edge between two different angle cut-offs (see Methods for more details on the measurement). The orientation domain areas are plotted as probability distributions (Fig. 3B). Using a Kolmogorov-Smirnov statistical test, we find that the distribution of domain sizes for 70 μM tubulin (n = 703) is distinct from the other distributions (p < 0.0001). Although it is clear that the data for 13.6 μM (n = 260) has a higher number of larger domains compared to 54.5 μM (n = 437), these distributions are not statistically distinct given the number of regions measured (p = 0.25).

We plot the mean domain size as a function of initial tubulin concentration to reveal a monotonically decreasing area with increasing tubulin concentration (Fig. 3Ci). Using the expected median lengths as a function of tubulin concentration (Fig. 2B), we can also plot the mean area size as a function of median microtubule length (Fig. 3Cii). We fit the mean area as a function of filament length to a linear function and a power law function for comparison (see log-log scale inset, Fig. 3Cii, fit parameters given in ESI Table 4). Both fits have two fit parameters, but the power-law fit has a lower Chi-squared value. The best fit power law has an exponent of 0.45 ± 0.03.

### Microtubule organization with crosslinkers

When crosslinkers were added to the system as the microtubules polymerize from dimers, they could bind laterally, and thus grow in two dimensions (Fig. 1A). We imaged microtubules polymerized in the presence of MAP65, an antiparallel crosslinker, to cause the condensation of new filament organizations (Fig. 1Bii). The MAP65 concentration was changed so that the percentage of tubulin dimers with a MAP65 crosslinker would range from 0% to 10% (Fig. 4, see ESI for details). We found that, as the crosslinker percentage was increased, the organizations changed from fan-like, splayed organizations to more condensed, smaller bundles with apparently pointed ends (Fig. 4). We were able to obtain the most data from the experiments with 13.6 μM tubulin, which clearly showed a transition from splayed (0%, 0.3%) to a co-existence between splayed and condensed bundles (1.5%, 3%), to all bundles (10%). Higher concentrations of tubulin also showed a transition from spayed fans to small, independent bundles as the crosslinker concentration increased (54.5 μM and 70 μM). Because these experiments had more tubulin and shorter microtubules, they necessarily had more filaments, making regions denser with bundles (Fig. 4).

**Figure 4.**
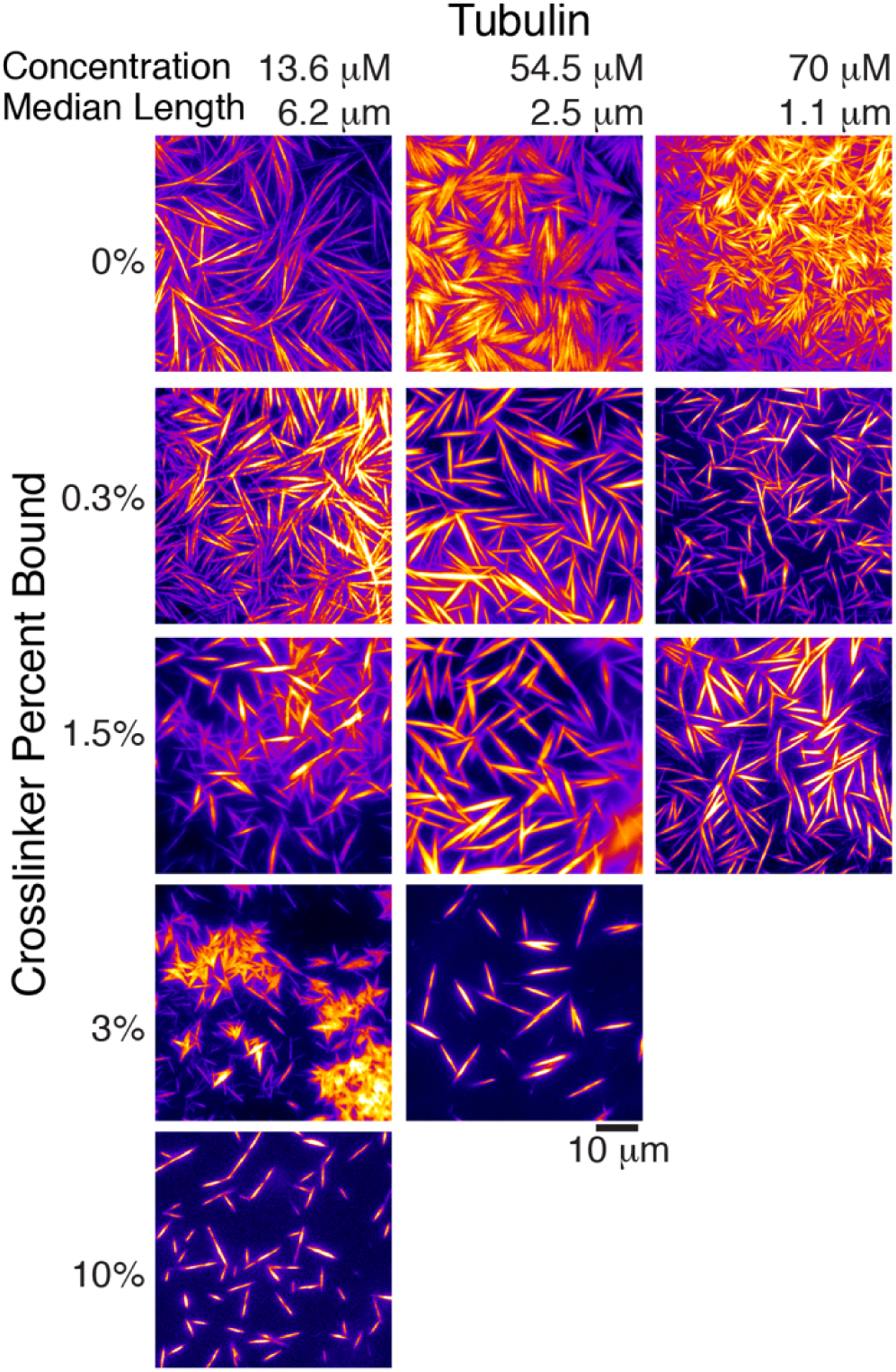
Phase diagram of microtubule self-organization final states. Microtubule patterns depended on the tubulin concentration, which controlled the filament median length, and the percent of MAP65 bound to the microtubules. Tubulin concentrations used were 13.6 μM (left column), 54.5 μM (middle column), and 70 μM (right column), corresponding to median filament lengths of 6.2 μm, 2.0 μm, and 1.1 μm, respectively. For each concentration of tubulin, a different concentration of crosslinking MAP65 resulted in the equivalent percent bound of MAP65 (see ESI for details). We tested MAP65 at 0%, 0.3%, 1.5%, 3%, and 10% for each tubulin concentration.

The condensed bundles are not long, thin bundles, as has been previously observed for Taxol-stabilized microtubules in the presence of MAP65 ^13,14^, or in prior active-matter experiments with kinesin crosslinkers ^7,30,31^. Instead, these bundles are finite in length and appear tapered on the ends. Indeed, they appear more like liquid crystal tactoids or mitotic spindles rather than traditional microtubule bundles ^26,32,33^. The definition of a tactoid is a spherical or spindle-shaped, birefringent droplet condensed from an isotropic domain. Prior work has suggested that the meiotic spindle is organized as a liquid crystal of short microtubule filaments condensed into the same tactoid shape ^8,34–38^. Most of that work was performed on meiotic spindles self-organized from Xenopus oocyte extracts ^39^.

### Microtubule tactoids

To test if the microtubule organizations we observe are tactoid-like in shape and birefringent character, we performed characterization tests. As suggested by the phase diagram, the condensation of these organizations appears to be driven by the presence of the MAP65 crosslinker. We tested if the crosslinker is incorporated into the bundle using two-color fluorescence imaging. The GFP-MAP65 appears to be bind everywhere the microtubules are present within the organization, including the tips of the tactoid (Fig. 5A).

**Figure 5.**
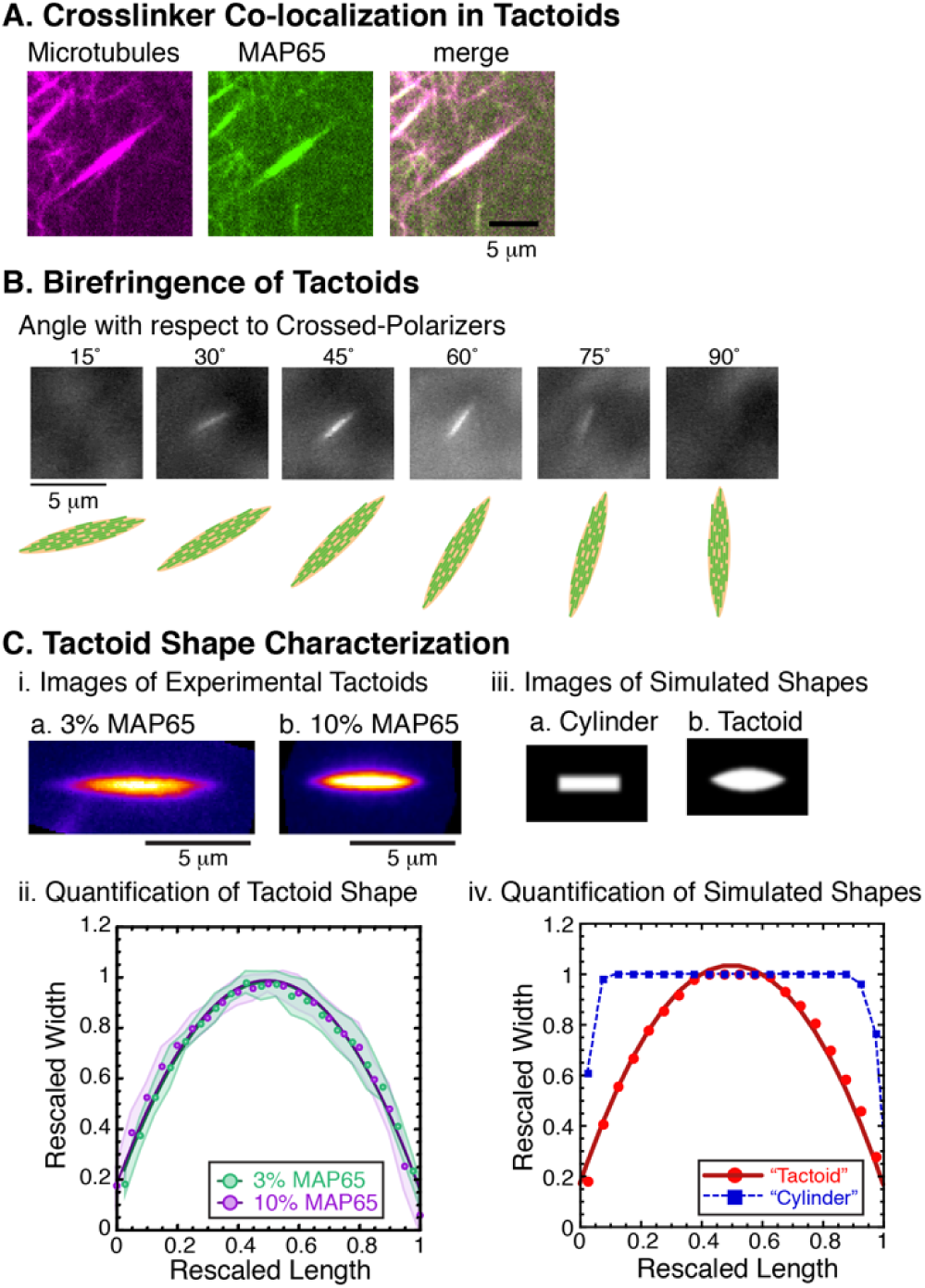
Tactoid characterization. (A) Same tactoid image in two color fluorescence to show the location of microtubules (left, magenta), GFP-MAP65 (center, green), and co-localization (merge). Scale bar is 5 µm. (B) Tactoids under crossed polarizers are rotated from 15° to 90°. Scale bar 5 µm. Cartoons depict the angle of the tactoid. (C) Tactoid shapes quantification using cross-sections perpendicular to the long axis to measure the width as a function along the length. (i) Representative images of tactoids for 13.6 µM tubulin in the presence of (a) 3% MAP65 or (b) 10% MAP65. (ii) Tactoid width rescaled by the maximum width and plotted as a function of the rescaled length for 3% MAP65 (green markers, n = 9), and 10% MAP65 (purple markers, n = 10). Shaded regions denote the standard deviation of the rescaled width. Data fit to Eq. 2 (green line for 3%, purple line for 10%) and best fit parameters given in ESI Table 6. (iii) Simulated bundle shapes to compare to tactoid shapes of (a) cylinder and (b) tactoid. (iv) Simulated shape width rescaled by the maximum width and plotted as a function of the rescaled length for the cylinder (blue squares) and the tactoid (red circles). Data for the tactoid shape is fit to Eq. 2 (red line).

Next, we test for birefringence of the microtubule organizations using cross-polarizers on a transmitted light microscopy microscope. This will demonstrate alignment and orientation of filaments within the organizations. We are able to rotate the sample to test if the intensity of light transmitted depended on the angle of the organization. Indeed, we find that the objects appeared birefringent, and allow the most light compared to the background when they are at a 45° angle with respect to the crossed polarizers (Fig. 5B).

Interestingly, we find that the birefringence appeared homogeneous along the length of the bundle structure, without dark regions at the ends. This result implies that the tactoid is a “homogeneous” type with poles located far away from the structure itself ^40^. This is different from bipolar tactoids where the poles are near the pointed edges. Homogeneous tactoid have the internal rods of the liquid crystal aligned parallel to the long axis, and not tilted at the points. We have depicted our cartoons (Fig. 5B, Fig 1A) to have the internal organization expected for homogeneous tactoids.

Finally, we quantified the shape of the tactoids for two different MAP65 relative concentrations (3% and 10%) and the same tubulin concentration (13.7 μM, Fig. 5Ci) using an through taking cross-sectional scans perpendicular to the long axis of the tactoid, rescaling the length to the maximum length and the width to the maximum width, averaging over multiple different tactoids, and fitting to a parabola of the form:

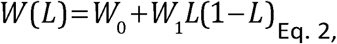

Where W(L) is the rescaled width as a function of the rescaled length (L) along the tactoid, W_0_ is the vertical offset of the parabola, and W_1_ is the amplitude of both the linear and squared terms of the parabola. When rescaled in this manner, we find that all the data for different tactoids and different experimental parameters overlay on top of each other and fit parameters are similar (Fig. 5Cii, ESI Table 6). Theoretically, the tactoids should have zero width at the two ends, but experimentally, this is not the case because the images are diffraction limited. Further, in order to average over tactoids of different lengths, we had to bin the data, which also resulted in a broadening of the tips. Together, these experimental aspects were accounted for using the offset, W_0_.

To test if the shape is truly a tactoid or a blunt-ended cylinder, we simulate two shapes using ImageJ with added Gaussian noise to act as diffraction blurring and perform the same cross-sectional analysis on these simulated shapes (Fig. 5Ciii, iv). Blurring of the data, similar to the resolution limit of the microscope, causes the edges of the cylindrical bundle to decrease gradually, but there is no mistaking the cylinder for a tactoid cross-section (Fig. 5Civ). The simulated tactoid clearly displays a parabolic shape in cross-section and fits to the same phenomenological equation as the experimental tactoids, including the offset, W_0_, due to the simulated diffraction and binning (Eq. 3, ESI Table 6). Thus, we conclude that the crosslinking drives microtubules to condense into tactoids which have a spindle-shape and are birefringent, as required.

### Characterization of tactoid aspect ratio

We expect that the cohesion strength between the filaments would affect the aspect ratio of the tactoid shape. We characterize the aspect ratio by measuring the length and width of the condensed bundles (Fig. 5A). We use the data set for 13.6 μM tubulin because it includes the most data points for different percentages of crosslinkers: 0.3%, 1.5%, 3%, and 10% MAP65 (Fig. 5A). For experiments with co-existence between tactoids and splayed fans, we measure the tactoids only. For each data set, we measure the length, width, and aspect ratio and plotted the probability distribution function (PDF, Fig. 5B).

As expected, the aspect ratio of the tactoids decreased with increasing adhesion strength (Fig. 6Ci). In this case, the adhesion is mediated by the percentage of MAP65 bound to the microtubules, and the aspect ratio decreases linearly with the log of the MAP65 percentage (fit information given in ESI Table 7). Typically, when aspect ratio of tactoids decreases, the width increases and the length decreases. Thus, we would expect for the width of the bundles to increase with increasing MAP65 percentage. We used the measurements of the widths and the lengths to determine if the width and length were both changing, or if one had more control over the aspect ratio. Surprisingly, the lengths of the tactoids decrease with increasing crosslinker percentage, but the widths are similar for all percentages (Fig. 5 B,C).

**Figure 6.**
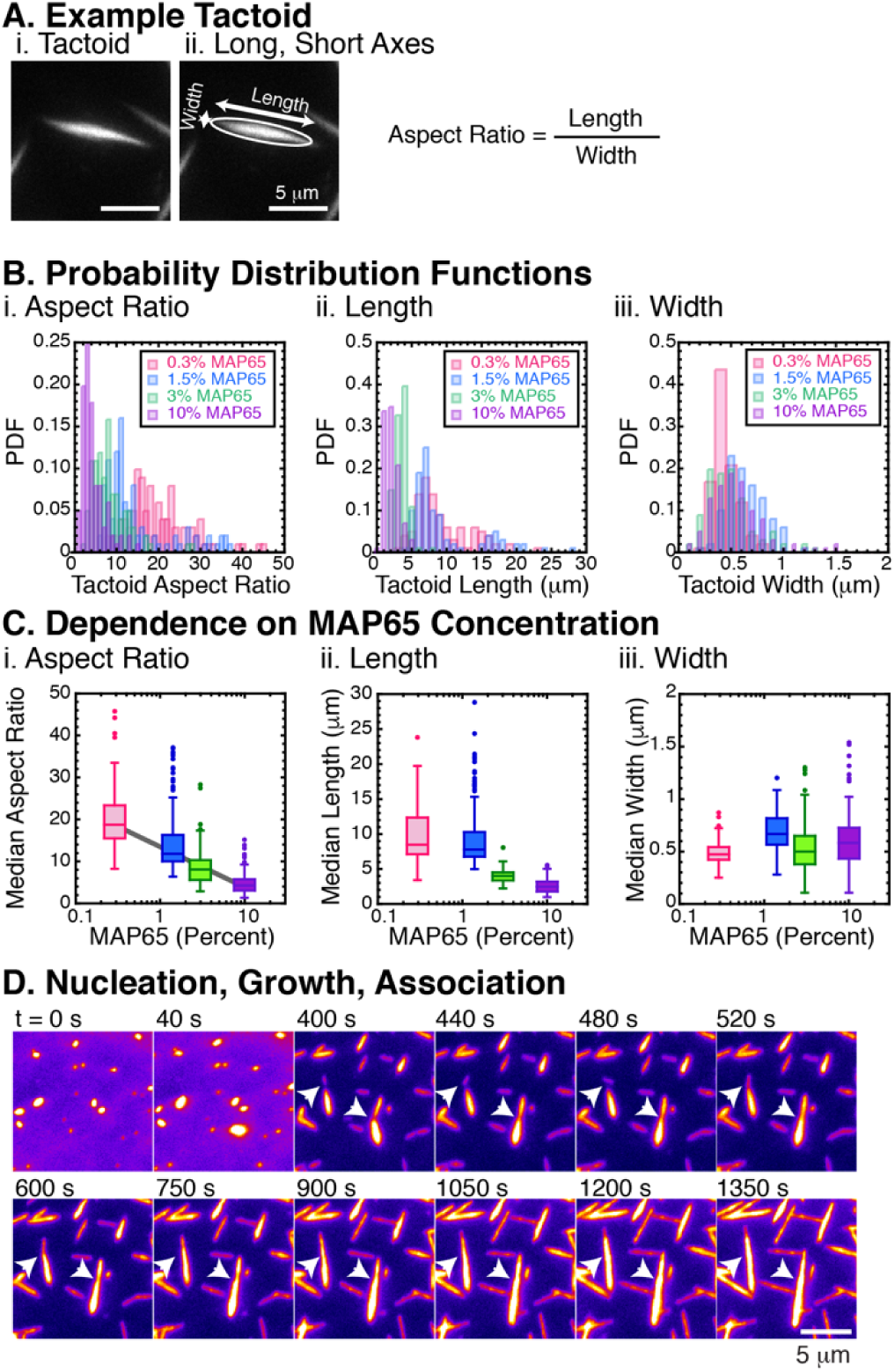
Tactoid characterization. (A) For regions of the phase diagram displaying tactoids, we measured the length, width, and computed the aspect ratio for >100 different tactoids. All data represented is from experiments with 13.6 μM tubulin. (B) Probability distributions (PDFs) of the (i) aspect ratio, (ii) length, and (iii) width for microtubule tactoids in the presence of 0.3% MAP65 (n = 101, pink), 1.5% MAP65 (n = 100, blue), 3% MAP65 (n = 101, green), and 10% MAP65 (n = 101, purple). (C) The median, quartiles, and outliers were computed for each distribution represented in (B) and plotted as box-whisker plots as a function of MAP65 percentage. (i) Tactoid aspect ratio has a logarithmic dependence on the MAP65 concentration; fit equation and best fit parameters are given (ESI Table S7). (ii) Tactoid length decreases as a function of MAP65. (iii) Tactoid width is insensitive to the MAP65 concentration. (D) Example time series of tactoid nucleation, growth, and association. Arrowheads indicate end-to-end association. Times are marked in seconds; scale bar 5 μm.

We also find that the median lengths of the tactoids for 10% MAP65 were shorter than the median length of the individual filaments when polymerized at 13.6 μM tubulin (compare Fig. 5Cii with Fig. 2B). We would expect that the length of the tactoid would be bounded at the lower end by the length of the filaments, which should be 6.2 μm. Yet, at 10% MAP65, the median tactoid length was 4.1 μm. Short tactoids could imply that the crosslinking MAP65 protein also acts as an additional nucleator of microtubules, nucleating filaments faster resulting in more, shorter filaments than we measured (Fig. 2). Indeed, Arabidopsis MAP65-1 has previously been shown to be a microtubule nucleator in addition to microtubule bundling activities ^41^.

### Tactoid assembly dynamics

In order to shed light on the mechanism of tactoid assembly and possibly explain why the widths are consistent with increasing adhesion between filaments, we directly observed the dynamics of the forming steady state system. At early times, small, anisotropic tactoids have already formed within the viewing area of the chamber (Fig. 6D). These individual puncta appear to have a spindle-shape and aspect ratio like tactoids, apparent as long as they are not smaller than the diffraction-limit. As these spindles grow, they do not diffuse laterally, but rather grow in place, mostly along their lengths (Fig. 6D). We observe two mechanisms by which the tactoids elongate. First, the spindles grow longer through polymerization of microtubules within the bundle (Fig. 6D). The bundles also get brighter over time, implying that microtubules are nucleating and growing within or attached to the sides of the growing tactoids (Fig. 6D).

A second mechanism that causes the bundles to grow long but not wide is through bundle annealing. We observed that if two tactoids are parallel and aligned to each other along their long axes, they can fuse end-to-end (Fig. 6D, arrows). Because the lateral diffusion of the tactoids is limited, two tactoids that intersect at an angle other than parallel (0 or 180°) often continue to grow independently, resulting in tactoids that appear fused at their ends or crossing. These tactoids do not diffuse into each other in our assays.

### Dynamics within tactoids

Our time series data showing tactoid growth and interaction (Fig. 1B, Fig 6D) imply that the spindle-like assemblies are not mobile. To directly probe the exchange of microtubules and MAP65 crosslinkers, we perform fluorescence recovery after photobleaching (FRAP) on the tactoids made with fluorescent tubulin and GFP-labeled MAP65 (Fig. 7). FRAP works by photobleaching the fluorescently-labeled proteins in half of the tactoid, and observing the fluorescence as it recovers due to the motion of the molecules within the assembly or exchange between the tactoid and the solution. It is used to determine the mobility of molecules in a region and can be used to determine the biochemical off and on rates for binding reactions ^42^. We can photobleach both the microtubules and MAP65 simultaneously and observe the recovery (Fig. 7A). We observe little to no recovery of the microtubule signal over the course of 30 minutes (Fig. 7Ai), but see significant recovery of the MAP65 signal even within 30 seconds (Fig. 7Aii).

**Figure 7.**
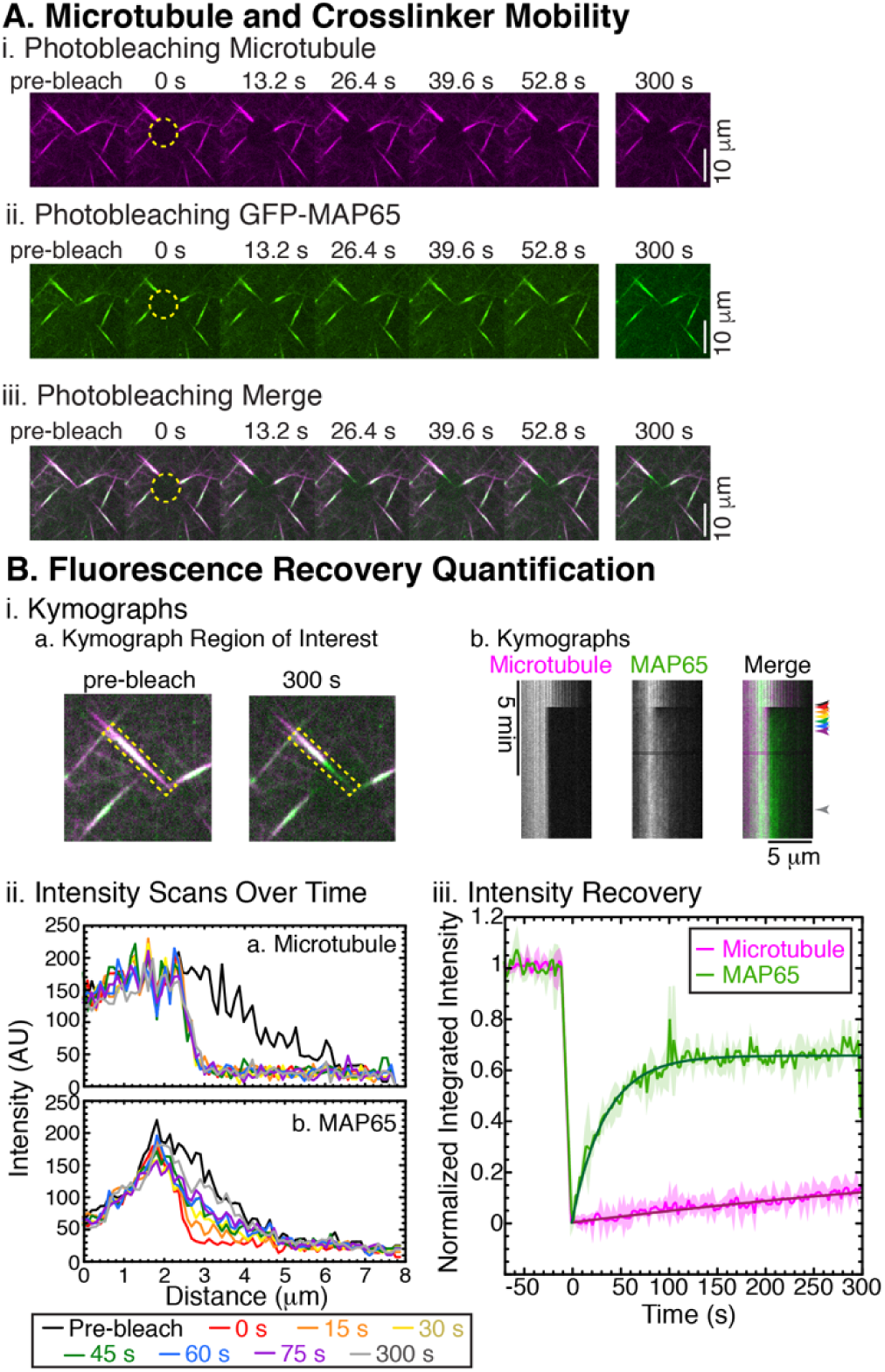
Tactoid Dynamics. (A) Fluorescence recovery after photobleaching of tactoids in two colors for (i) microtubules (magenta) and (ii) GFP-MAP65 (green) with (iii) merged images. Dashed circle indicates bleached area at t = 0 s. Times as indicated at 13.2 s intervals; last frame at 300 s; scale bar 5 μm. (B) Quantification of photobleaching. (i) Kymographs show fluorescence recovery. (a) Linear region, 3 pixels wide (yellow dashed outline) of selection overlaying the tactoid with microtubules (magenta) and GFP-MAP65 (green) merged shown (left) pre-bleach and (right) 300 s after bleach. (b) Kymograph of tactoid FRAP showing (left) microtubules, (middle) GFP-MAP65, and (right) merged image of microtubules (magenta) and GFP-MAP65 (green). (ii) Intensity along the tactoid length for the (a) microtubule and (b) MAP65 at different times: pre-bleach (black line), 0 s (red line), 15 s (orange line), 30 s (yellow line), 45 s (green line), 60 s (blue line), 75 s (purple line), and 300 s (gray line). Time points denoted with same-colored arrowheads next to the kymographs in (above). (iii) The fluorescence recovery over the entire bleached region was quantified to demonstrate that the GFP-MAP65 (green line) recovers to about 60% of the initial intensity, while the microtubules (magenta line) regain a minimal level (<20%). The shaded regions denote the standard deviation for two separate bleaching experiments on different areas.

We directly demonstrate the recovery using kymographs (space-time representations) by taking a region of interest along a tactoid (Fig. 7Bia) and sequentially displaying the images side-by-side in the vertical direction (Fig. 7Bib). It is clear in the kymograph that the intensity of the microtubule does not recover, while the intensity of the MAP65 recovers (Fig. 7Bib). Using the kymographs, we can plot intensity profiles along the tactoid for different time points (colored arrowheads denote time points in Fig. 7Bii). The microtubule intensity scans do not recover over 300 s (Fig. 7Biia), while the GFP-MAP65 gradually recovers over time and displays a slightly reduced intensity at 300 s compared to pre-bleaching (Fig. 7Biib).

The fact that the MAP65 recovers implies that it is highly mobile within the tactoid structure. The pattern of recovery occurs all along the length, and does not appear to move from the unbleached side into the bleached region (Fig. 7Biib) implying that the exchange of MAP65 is likely coming from free solution and not strictly from MAP65 molecules within the tactoid. Additionally, the fact that MAP65 is capable of recovery implies that the microtubules of the tactoid, although bleached, are not damaged. It is perhaps not surprising that the recovery of the MAP65 signal is fast given its known low binding constant (K_D_ ∼ 1.2 µM), and previously measured fast turn over rates ^13,19^.

The lack of recovery of the microtubule signal implies that the microtubules are not mobile within the tactoid on the timescale of the experiment. It is likley that the microtubules are jammed or arrested in this state by the crosslinker molecules. The limited mobility of microtubules within the patterns may be due to the strength of the crosslinker. Perhaps the MAP65 has a strong affinity compared to the thermal fluctuations. We have previously shown that MAP65 is capable of overpowering kinesin-1 motors in a microtubule filament gliding assay ^13,14^. Kinesin-1 is an incredibly powerful molecular motor, each being able to create 5 pN of force over 8 nm, the equivalent of roughly 10 k_B_T of energy. In a gliding assay configuration, where thousands of motors can be simultaneously engaged to propel the microtubules, the maximal energy that can be exerted is 10^4^ k_B_ T. Yet, MAP65 is capable of stalling these motors when crosslinking two microtubules in opposing directions. In order to loosen the interaction between the filaments, we lowered the MAP65 concentration to 0.3%, yet we still see no recovery of the spindle-like assemblies. Further, when there is no MAP65 present, photobleaching recovery is also minimal, implying that the system becomes arrested regardless of the presence of crosslinker.

Our results imply that the spindle-like bundles we have created do not display liquid-like properties as liquid crystals, meiotic spindles, or actin tactoids have been shown to do ^25,26,31,43,44^. This may be due to the crowding agent, methylcellulose, we employed. Methylcellulose is a large, viscous polymer that can form bundles on its own at high concentrations ^45^. Here, we use it at 0.3% in the chamber, which should be in the isotropic network phase ^45^. The version we use is large (88 kD) and viscous (4000 cP). The concentration we use (0.3%) is below the overlap concentration and methylcellulose is a negatively-charged polymer, so our polymers are likely individual polymer chains with a radius of gyration approximately 30 nm. The methylcellulose is probably acting only as a viscosity agent and depletion agent to slow and bundle microtubules, but we do not know this exactly. Future studies using other crowding agents such as PEG or dextran at various molecular weights and different viscosities could help determine if the crowding agent is responsible for the immobility of our spindle-like assemblies.

Our FRAP measurements also show that individual filaments have low mobility. It is possible that the methylcellulose or the surface coating polymers (Pluronic F127) are limiting the mobility of individual microtubule filaments within the organizations. This would be surprising, because we would expect that the crowding agent would be excluded from the organizations. Future work with fluorescent crowding agents to measure the dynamics of the crowder would illuminate the mobility of the network of methylcellulose or other crowding agents employed.

## CONCLUSION

We have demonstrated that the active polymerization of microtubules from tubulin dimers can lead to beautiful and biologically relevant self-organized structures in a minimal experimental system (Fig. 1). Microtubules are rigid rods, that are longer than they are wide, so they should naturally align into nematic organization at high density. In this study, we have added a crowding agent, methylcellulose, to increase the local concentration of filaments and crowd the system to the surface (quasi-2D). Our minimal system with crowding alone can create fan-like organizations of polymerizing microtubules, displaying local order, as expected, with ordered domains of limited size that scale with filament median length (Fig. 3, 4). The mechanism is from initial filaments with different orientations all nucleate and grow the local domains simultaneously, causing grain boundaries to form where these domains meet. The size of the domains is controlled by the length of the microtubules in the organization, which is in turn controlled by the concentration of tubulin used (Fig. 2).

When crosslinking proteins are employed in the networks, the microtubules align and condense into bundles that resemble tactoids or mitotic spindles (Figs. 4-7). These assemblies are likely homogeneous tactoids since they are birefringent along their entire length and have a tactoid shape that is longer and thinner (Fig. 5). Their aspect ratio scales with the adhesion between the filaments, as expected, although the width does not change significantly – only the length (Fig. 6). Using FRAP, we show that the microtubules within tactoids are not truly “liquid-like” as they do not appear to have the mobility required for fluorescence recovery (Fig. 7). The MAP65, on the other hand, is highly dynamic, and likely exchanging with the free MAP65 in solution (Fig. 7). It is interesting that a highly dynamic, and “weak” interacting crosslinker can so tightly glue together the microtubules. This is because the dynamic equilibrium enables a lot of MAP65 to bind, even as individuals exchange with solution. Future work using different crowding agents, concentrations, and molecular weights may determine if the crowding agent is the cause for the limited mobility and fixed width of the assemblies.

Our new experimental system is an exciting leap forward for self-organization of spindle-like assemblies. With very few components, we have recreated assemblies with the proper shape to be a model *in vitr*o spindle. Further, we demonstrate that the motor property of active crosslinkers needed for the assembly of cellular spindles, such as Eg5, is likely not the driving force for the organization into the spindle shape. Rather, our results demonstrate that the crosslinking activity of these motors and other crosslinkers are likely needed to form the spindle shape.

Our bottom-up approach is important to understand the physical principles behind the exciting, complex, and dynamic organizations observed in live cells. The minimally-reconstituted system we demonstrate here is not a perfect model of the spindle due to its longer, homogeneous tactoid shape. The long shape is likely due to the long filaments (≥1 µm) we employed. In true spindles, the length of the microtubules can be dynamically controlled and adjusted even within the structure, as demonstrated in high resolution electron microscopy studies of serial sections along spindles ^46,47^. Prior simulations of the meiotic spindle that used longer filaments also had the same long, thin shape that we see in this study ^48^.

Another difference between our tactoid assemblies and true spindles is the internal reorganization and dynamics of the filaments. Meiotic spindles incredibly liquid-like with fast turn-over of filaments and even the ability to coalesce when two come close together. Conversely, many cellular mitotic spindles have a mid-zone that turns over and appears liquid-like, and an end-zone that is jammed or arrested to hold the organization together. Our work lays the groundwork to recreating more realistic, yet minimal, assemblies to test biophysical and biochemical hypotheses of mitotic spindle organization, dynamics, and complexity.

Recent work on actin has also shown that short actin filaments can create true, liquid-like tactoids in the presence of weak crosslinkers ^26^. These are exciting structures, but less biologically relevant, since actin does not form spindles in cells – only microtubules form spindles in the process of cell division. Other work using microtubules were also able to recapitulate spindle-like organizations by confining microtubules grown within an emulsion droplet from purified microtubule-organizing centers from cells ^49^. In this system, the spindle size was set by the size of the droplet because the filaments spanned the droplet and buckled, and required a microtubule nucleation complex. Meiotic spindles set their own intrinsic length, possibility by controlling the length of the microtubules. Indeed, recent work has shown evidence for microtubule length regulating proteins, such as microtubule severing enzymes, to control spindle length ^48^.

Future work utilizing this system could overcome the immobility by specifically adding in active motor proteins, such as kinesin-1, into the system. The kinesin-1 could be tetramerized as previously performed for active liquid crystals or bound to the surface in a gliding configuration. Other kinesin or dynein family microtubule-based motor proteins could also display different activities, since each has different binding and force generation properties. Furthermore, as we have used MAP65 as a relatively strong crosslinker, and future works could employ another weaker, homologous analog such as PRC1 or Ase1.

## Acknowledgements

We thank Shuang Zhou for use of his polarization microscope. BE, CFdR, and JLR were funded by NSF Inspire Award #1344203 to JLR. SS was funded by MURI grant #67455-CH-MUR to S. Thayumanavan. BE was also partially funded by NSF MRSEC grant # 1420382 to Seth Fraden at Brandeis University. We would like to acknowledge the help and guidance of Prof. Bruce Goode (Brandeis University) for mentoring BE in the summer of 2018. The authors would like to acknowledge the help of Lena Herbst, Austin Morrisey, and Talia O’Shea for their helpful feedback in lab. We thank the University of Massachusetts Institute for Applied Life Sciences (IALS) light microscopy core for use of the spinning disc and FRAP system for use in this work.

