## Supplemental Data for "Self-Organization of Spindle-Like Microtubule Structures"

Electronic Supplemental Information for
**Self-Organization of Spindle-Like Microtubule Architectures**

Bianca Edozie[1,2], Sumon Sahu[1], Miranda Pitta[3], Carline Fermino do Rosario[1,4], Jennifer L. Ross[1,4#]

1. Department of Physics, University of Massachusetts Amherst, MA 01003
2. Department of Biochemistry and Molecular Biology, University of Massachusetts Amherst, MA 01003
3. Hopkins Academy, Hadley, MA 01035
4. Molecular and Cellular Biology Graduate Program, University of Massachusetts Amherst, MA 01003

[#]corresponding author

**Detailed Experimental Methods.**

*MAP65 concentration calculations*. The concentration of MAP65 added to each sample depended on the final percentage of tubulin dimers we wanted occupied by the MAP65, the concentration of tubulin dimers, and the equilibrium binding constant for the MAP65 to microtubules, 1.2 μM {Talin 2012}. The chemical reaction is assumed to be a simple binding reaction with one-to-one stoichiometry between tubulin and MAP65:

$$[TUB_{free}] + [MAP_{free}] \rightleftharpoons [MAP - TUB], \qquad \text{Rxn 1}$$

Where $[TUB_{free}]$ is the concentration of tubulin in microtubule form without a MAP65 bound, the $[MAP_{free}]$ is the concentration of MAP65 in solution that is not bound to tubulin and $[MAP - TUB]$ is the concentration of the tubulins or the MAP65 molecules that are complexed. Since the reaction is assumed to be one-to-one, this is the same concentration for either MAP65 or tubulin.

The general differential equation to describe the reaction is:

$$\frac{d[MAP - TUB]}{dt} = k_{on}[TUB_{free}][MAP_{free}] - k_{off}[MAP - TUB], \qquad \text{Eqn 1}$$

Where $k_{on}$ is the on-rate for MAP65 to bind to tubulin with units of per molar per time, and $k_{off}$ is the off-rate of MAP65 dissociating from tubulin with units of per time. At equilibrium, amount MAP65 binding and unbinding is equal, so the concentration of $[MAP - TUB]$ will be constant over time. Thus, the left hand side of the equation will equal zero, and we can solve for a single equilibrium dissociation constant:

$$K_D = \frac{k_{off}}{k_{on}}, \qquad \text{Eqn 2}$$

Resulting in:

$$K_D = \frac{[MAP_{free}][TUB_{free}]}{[MAP - TUB]}, \qquad \text{Eqn 3}$$

We want the percentage of tubulin dimers with a MAP65 bound. The percent bound is equal to:

$$\%Bound = 100 * \frac{[MAP - TUB]}{[MAP - TUB] + [TUB_{free}]}, \qquad \text{Eqn 4}$$

The sum of the $[MAP - TUB]$ and the $[TUB_{free}]$ equals the total tubulin concentration added to the chamber:

$$[TUB_{total}] = [MAP - TUB] + [TUB_{free}], \qquad \text{Eqn 5}$$

This is a parameter we can control, and equals 13.6 μM, 54.5 μM, or 70 μM, depending on the experiment. Combining equations 4 and 5, we can find that:

$$[MAP - TUB] = (\%Bound)([TUB_{total}]), \qquad \text{Eqn 6}$$

In equation 3, we now know the left hand side, $K_D$, the denominator on the right hand side, $[MAP - TUB]$, and the parameter $[TUB_{free}]$. We want to know the amount of MAP65 to add to the chamber, $[MAP_{total}]$, to get the specified $\%Bound$, but equation 3 can only give the amount of MAP65 free in solution, $[MAP_{free}]$.

Using a spreadsheet, we can calculate $[MAP - TUB]$ from equation 6 and $[MAP_{free}]$ from equation 3 combined with equation 5. We can add these together to determine the concentration of MAP65 to add to the chamber, total:

$$[MAP_{total}] = [MAP_{free}] + [MAP - TUB], \qquad \text{Eqn 7}$$

Alternatively, you can solve for $[MAP_{total}]$ in one step using this equation:

$$[MAP_{total}] = \%Bound \left( \frac{K_D}{1 - \%Bound} + [TUB_{total}] \right), \qquad \text{Eqn 8}$$

Using this method, we altered the final concentration added to the chamber with the tubulin as given in Table 1, below.

**ESI Table 1.** Table of the concentrations of MAP65 used in this study depending on the tubulin concentration and desired percent bound. $K_D$ = 1.2 μM.

| | Tubulin Concentration | | |
|---|---|---|---|
| Percent MAP65 Bound | 13.6 μM | 54.5 μM | 70 μM |
| 0 | 0 | 0 | 0 |
| 0.3% | 0.0044 | 0.017 | 0.021 |
| 1.5% | 0.22 | 0.84 | 1.07 |
| 3% | 0.45 | 1.67 | 2.13 |
| 10% | 1.49 | 5.58 | 7.13 |
| | MAP65 Concentration (μM) | | |

**ESI Table 2.** Fit equations for cumulative probability distributions of microtubule lengths (Fig. 2A).
Fit equation:

$$CDF = 0.5 + 0.5 erf\left(\frac{Ln(L_C) - \mu}{\sqrt{2\sigma}}\right)$$

| Tubulin Concentration | μ | σ | $\chi^2$ |
|---|---|---|---|
| 13.6 μM GMPCPP | 1.8286 ± 0.0008 | 0.260 ± 0.001 | 0.0519 |
| 15.5 μM GMPCPP | 1.766 ± 0.002 | 0.505 ± 0.003 | 0.0457 |
| 30 μM GMPCPP | 1.632 ± 0.002 | 0.280 ± 0.003 | 0.107 |
| 46.5 μM GMPCPP | 0.962 ± 0.003 | 0.497 ± 0.005 | 0.0305 |
| 54.5 μM GMPCPP | 0.600 ± 0.002 | 0.367 ± 0.003 | 0.0414 |
| 54.5 μM GMPCPP | 1.2387 ± 0.0009 | 0.333 ± 0.001 | 0.0210 |
| 45.5 μM GTP | 2.512 ± 0.001 | 0.512 ± 0.002 | 0.0588 |

**ESI Table 3.** Linear fit equation for microtubule length as a function of tubulin concentration (Fig. 2B).
Fit equation:

$$L_C = L_0 - m[TUB]$$

| Data | $L_0$ | m | $R^2$ |
|---|---|---|---|
| CDF, GMPCPP | 7.4 ± 0.7 | 0.09 ± 0.02 | 0.88195 |

**ESI Table 4.** Linear and power law fits for mean orientation domain areas as a function of microtubule contour length (Fig. 3Cii).
Linear fit equation:

$$A(L_C) = A_0 + mL_C$$

Power Law fit equation:

$$A(L_C) = A(L_C)^{\gamma}$$

| Fit Equation | | | | |
|---|---|---|---|---|
| **Linear** | **$A_0$** | **m** | **$R^2$** | $\chi^2$ |
| | 24 ± 5 | 7 ± 1 | 0.968 | 20.1 |
| **Power Law** | **A** | $\gamma$ | | $\chi^2$ |
| | 28 ± 1 | 0.45 ± 0.03 | | 2.6 |

**ESI Table 6.** Tactoid shape equation for 3%, 10% MAP65 and simulated tactoid (Fig. 5C, ii, iv)
Fit equation:

$$W(L) = W_0 + W_1 L(1 - L)$$

| **Data Set** | **$W_0$** | **$W_1$** | $\chi^2$ |
|---|---|---|---|
| 3% MAP65 | 0.15 ± 0.01 | 3.31 ± 0.06 | 0.012 |
| 10% MAP65 | 0.17 ± 0.02 | 3.3 ± 0.1 | 0.034 |
| Simulated Tactoid | 0.12 ± 0.01 | 3.46 ± 0.07 | 0.016 |

**ESI Table 7.** Aspect Ratio as a function of MAP65 percent bound (Fig. 6Ci).
Fit equation:

$$R = R_0 - q log[\%MAP65]$$

| **$R_0$** | **q** | **$R^2$** |
|---|---|---|
| 13.5 ± 0.4 | 9.7 ± 0.6 | 0.99 |

**ESI Table 8.** Fluorescence recovery after photobleaching (FRAP) intensity recovery data (Fig. 7Ciii).
Fit equation:

$$I(t) = I_{\infty}\left(1 - exp(-t/\tau)\right)$$

| | $I_{\infty}$ | $\tau$ | **$R^2$** | $\chi^2$ |
|---|---|---|---|---|
| Microtubule | 0.4 ± 0.1 | 700 ± 300 | 0.86 | 0.025 |
| GFP-MAP65 | 0.654 ± 0.004 | 34 ± 1 | 0.93 | 0.193 |
